# Honeyweed (*Leonurus sibiricus*) and Buckwheat (*Fagopyrum esculentum*) supplemented diet improve growth performance, lipid profile and hematological values of broiler chicks

**DOI:** 10.1101/2020.06.18.159764

**Authors:** M.T. Abedin, S.T. Rahman, M.M. Islam, S. Hayder, M.A. Ashraf, J. Pelc, M.A. Sayed

## Abstract

The study was conducted to investigate the combined effects of honeyweed (HW) and buckwheat (BW) supplemented diets on the growth, feed intake, serum lipid profile and blood parameters in broiler chicks. One hundred fifty (150) day old chicks (Cobb-500) were divided into five groups *viz*. T1 (Commercial control, CC), T2 (FF+10% BW), T3 (FF+10% BW +5% (w/v) HW powder), T4 (FF+10% BW +10% (w/v) HW powder), T5 (FF+10% BW +15% (w/v) HW powder) in complete randomized design with five (5) replications, each of which contain six (6) birds. The CC feed and the FF with HW and BW supplemented diet fed on broiler for 30 days. At the end of the study, the body weight gain, FCR and mortality rate were calculated. It was found that T3 (FF+10% BW +5% (w/v) HW powder) diet significantly (p <0.05) showed the good FCR, mortality rate and body weight gain. Interestingly, T3 decreased the serum cholesterol, triglycerides (TG), low density lipoprotein (LDL), increased high density lipoprotein (HDL) and improves blood parameters significantly (p < 0.05). Our results suggested that this newly formulated feed T3 (FF+10% BW +5% (w/v) HW powder) could be considered as an alternative natural feed additive to hazardous synthetic antibiotics.

## Introduction

In the 19^th^ century researchers are emphasized on increasing the food production but in the 20^th^ century the whole world is intensified to quality and secured food rather than quantity. Poultry industry plays an important role to meet the demand of ever increasing world population. Poultry meat is one of the easiest and cheapest sources of protein. Quality poultry meat production could be increased through poultry rearing but the unavailability of quality feed and the disease susceptibility are the major challenges for the poultry industry. Antibiotics represent an extremely important tool in the efficient production of meat and eggs by improving growth rate, reproductive performance, feed utilization and reducing mortality. It is not only improved poultry performance but also it has some economic benefits. Unfortunately, the long term and extensive use of antibiotics result in selection for the survival of resistant bacteria strain (Suresh *et al*., 2018). At the same time the dissemination of antibiotic resistant strains of pathogenic and non-pathogenic micro-organisms into the environment and their further transmission to human through the food chain could be lead to serious consequences on public health (Founou *et al*., 2016). The great number of antimicrobial residues was found in livers, kidneys, thigh and breast muscles in poultry meat which is now the major alarming issue for health safety (Sattar *et al*., 2014). There are some recently published data indicated that antibiotic residues make consumers, particularly children and the elderly, cause to liver and kidney diseases. These may have carcinogenicity and genotoxicity effects on human health as well (Lee *et al*., 2001). To upsurge the profitability in the poultry industry, farmers have been used antibiotics even for one week long without any prescriptions by the doctors. It has been studied that around 75%, 90% and sometimes 50–100% veterinary antibiotic being excreted through the feces and urine without absorption by the digestive system (Kim *et al*., 2011). These feces have been used as poultry manure in the crop field to improve the soil health. Hence, the antibiotic residue create adverse effects and destroy the growth of beneficial microorganism in the soil. In addition, residue can be uptaken by plants and accumulated in food grains. Subsequently, antibiotic resistant bacteria can transmit directly and indirectly through the food chains, air, water and soil (Grenni *et al*, 2018). The antibiotics had significant negative effect on soil microbes and destroy the soil properties (Kong *et al*, 2006). Elsewhere integrated fish farming combines livestock production with fish farming is very much popular in South East Asia. There are many poultry shed made directly on a fish pond for supplying as fertilizer and supports the growth of photosynthetic organisms. This system favor antimicrobial-resistant bacteria in the aquatic environment (Kong *et al*., 2006). However, many countries have already taken action to reduce the use of antibiotics in food-producing animals including Bangladesh. For example, since 2006, the European Union has banned the use of antibiotics for growth promotion. Consumers are also driving the demand for meat raised without routine use of antibiotics, with some major food chains adopting “antibiotic-free” policies for their meat supplies. Lot of alternatives have been proposed and examined in poultry production to reduce this problem. Among these, phytogenic feed additives (PFAs) are natural bioactive compounds, derived from plants and incorporated into animal feed to enhance productivity draw major attention (Karásková *et al*., 2015). From the previous studies it is proved that phytogenic feed additives has antimicrobial effect, enhance digestibility and improve performance in poultry production. (Sayed *et al*., 2015; Islam *et al*., 2016).

An herbaceous annual or biennial medicinal herb, honeyweed (*Leonurus sibiricus*), could be the alternative to the antibiotic. Due to its bioactive compound such as diterpenes, triterpenes, flavonoids, sterols and phenolic acids, it is effective for the treatment of common diseases like diabetes, cancer, kidney and heart diseases (Sayed *et al*., 2016). Recently the cardiovascular protection effect of leonurine has drawn most attention and many other activities have been assigned to various compounds from the genus *Leonurus* (Zhang *et al*., 2018) and it has anti-microbial activity (Chua *et al*., 2017).

In addition that buckwheat (*Fagopyrum esculentum*) is a pseudo cereal could also be the alternative synthetic feed additive. It is a dicotyledonous crop under the family of polygonaceae with higher amounts of all essential amino acids such as lysine, arginine and aspartic acid makes it unique and superior protein sources as well from other cereals (Zhang *et al*., 2012). Buckwheat consist of some important bioactive compounds that can play an antioxidant, anti-inflammatory and anti-hypertensive role due to the presence of flavones, flavonoids, phytosterols and myo-inositol (Zhang et al., 2012). Rutin is a flavonol glycoside which helps to strengthen and increase flexibility in blood vessels was not found in cereals and pseudocereals except buckwheat (Watanabe, 1998). Buckwheat is rich in unsaturated fatty acids which helps in human disorders such a cardiovascular diseases, hypercholesterolemia or hypertension. The addition of protein products of buckwheat to diets significantly lowers the blood cholesterol level, particularly that of low density lipoproteins (LDL) and very low density lipoproteins (VLDL) (Tomotake *et al*., 2006). The antimicrobial effect of buckwheat has also been report (Świątecka et al., 2013).

In previous study, it was found that supplementation of buckwheat seed powder in broiler feed significantly improve growth performance, decrease mortality rate, increase serum high density lipoproteins (HDL-cholesterol) and reduce serum triglycerides (Sayed *et al*., 2016). Interestingly, literature review suggested that the combined effect of buckwheat and honeyweed was not observed yet. Considering the medicinal advantages of buckwheat seed and honeyweed, the current study was designed to evaluate the growth performance and serum biochemical metabolites in broiler chicks.

## Materials and Methods

The present study was conducted in a commercial poultry farm (Joynal Agro and Feed) Pakerhat, Khansama, Dinajpur, in Bangladesh. The animal experiment was conducted by following the rules of intuitional animal care and use committee to ensure the ethical treatment. Biochemical parameters were measured in the Laboratory, Department of Biochemistry and Molecular Biology, Hajee Mohammad Danesh Science and Technology University (HSTU), Dinajpur.

### Collection and preparation of buckwheat and honeyweed

Buckwheat (BW) was purchased from the local market of Panchagar and honeyweed was collected from road site at Ataikula (24.0313° N, 89.4068° E), Pabna, Bangladesh. The honeyweed leaves were washed with running tab water and dried at room temperature. Dried leaves were kept in the oven at 60°C for 72 hours. Subsequently, oven dried leaves were powdered by grinder. Powders were fed by making solutions of 5, 10 and 15% with water.

### Feed formulation

Sundried and grinded, buckwheat, corns, meat and bone meal, rice polish, soybean meal, soybean oil and other feed ingredients were purchased from local market of Dinajpur, Bangladesh and then directly mixed with manually prepared diets to meet the nutrient requirement of broiler chicks (Table-1) provided by the National Research Council (NRC, 1994). Vitamin premix was purchased from Renata Animal Health Ltd. Two types of feed were formulated for broiler starter (1-15 days) and broiler grower (16-30 days) for conducting the experiment.

**Table 1:** Composition (%) of experimental diet in different rearing period

| **Diet Composition** | **1-15days (Starter)** | **16-30days (Grower)** |
| --- | --- | --- |
| Maize | 40.5 | 43.4 |
| Soyabean meal | 24.36 | 21.57 |
| Rice Polish | 10 | 10 |
| Meat and Bone meal | 4 | 4 |
| Protein Concentrate | 5.5 | 5.5 |
| Soyabean oil | 3 | 3 |
| Lime stone | 0.5 | 0.5 |
| DCP | 1 | 1 |
| Salt | 0.3 | 0.3 |
| Methionine | 0.12 | 0.1 |
| Broiler Premix* | 0.31 | 0.25 |
| Toxin binder | 0.3 | 0.3 |
| Coccidiostates | 0.05 | 0.02 |
| Lysine | 0.01 | 0.01 |
| Enzyme | 0.05 | 0.05 |
| Buckwheat | 10 | 10 |
| **Chemical composition of calculated nutrient /Kg feed** |  |  |
| Metaboisable energy (kcal/kg) | 3079.73 | 3163.61 |
| Crude Protein | 23.00 | 22.47 |
| Crude Fiber | 3.84 | 3.71 |
| Calcium | 1.01 | 0.86 |
| Available phosphorus | 0.51 | 0.56 |
| Methionine | 0.47 | 0.42 |
| Lysine | 1.38 | 1.15 |

\*Added broiler premix (Renata Animal Health Ltd.) contained: vitamin A: 4800 IU; vitamin D: 960 IU; vitamin E: 9.2 mg; vitamin K3: 800 mg; vitamin B1: 600 mg; vitamin B2: 2 mg; vitamin B3: 12 mg; vitamin B5: 3.2 mg; vitamin B6: 1.8 mg; vitamin B9: 2 mg; vitamin B12: 0.004 mg; Co: 0.3 mg; Cu: 2.6 mg; Fe: 9.6 mg; I: 0.6 mg; Mn: 19.2 mg; Zn: 16 mg; Se: 0.48 mg; DL-methionine: 20 mg; L- lysine:12 mg. Enzyme: phytase (Renata Animal Health Ltd. Bangladesh). Experiment. They were vaccinated against Newcastle and Gumboro diseases four times at one week interval.

### Experimental design and raring the birds

The animal experiment was conducted as described previously (Siddiqui *et al*., 2015). In brief, a total 150 one-day old broiler chicks (Cobb-500) were purchased from Aman Chicks Ltd. Birds were reared at brooding house to adjust with the environmental condition up to 10 days. Feeds and water were provided to the broiler chick *ad labitum*. Electric bulb was used to maintain temperature in the brooder house. Throughout the study, standard temperature was maintained and temperature gradually decreasing from 32 °C to 24 °C. After 10 days, they were randomly divided into five dietary treatment groups of 30 chicks each; each treatment was composed of five replicates with six birds in each in a complete randomized design. The birds were housed on floor and routinely managed. All the birds were examined twice daily for abnormal clinical signs (restlessness, lordosis, abnormal gait, vices and depression) as well as feed intake throughout the experiment. They were vaccinated against Newcastle and Gumboro diseases. During the 30 days of experimental period, growth performances of the birds were recorded. Before giving dietary supplementation treatment, body weight (g) was recorded for each group of birds. Then body weight (g) and feed consumption were recorded daily and body weight gain (g) and feed conversion were then calculated. Mortality (%) was recorded throughout the study. Feed consumption is the amount of feed consumed every day. It was calculated for each treatment at daily basis. At the end of the week, the residual amount of feed was weight and subtracted from the known weight of feed at the beginning of week. The product was divided by the total number of birds.

### Collection of blood, determination of lipid profile and blood parameters

The blood was collected from experimental and control birds at 30^th^ day by using a sterilized disposable syringe and needle by wing vein puncture without using any anticoagulant. Each of the syringes with blood sample was kept at normal temperature in an inclined position. After 20 minutes, the serum was collected and centrifuged for 15 minutes at 2500 rpm. After centrifugation, the supernatant were carefully collected by a micropipette and preserved in eppendorf vials. The collected serum samples were used for the determination of total cholesterol, HDL-cholesterol, LDL-cholesterol, triacylglycerols by using the kits purchased from CRESCENT diagnostic, Jeddah, K.S.A. Haematological paprameter were determined the automatic blood parameter analyzer (Pentra ES 60).

### Statistical Analysis

All data were analysed with the IBM SPSS statistics 22. A P-value of <0.05 was considered to indicate a significant difference between groups, and a comparison of means was made using Duncan’s multiple range test (Steel *et al*., 1984). The results are expressed as average ± standard deviation of five replications (n = 5). Data points bearing different letters are significantly different. (P<0.05)

## Results

### Growth performance

The effect of each formulated diet on growth performances of broiler chicks (Cobb-500) are shown in table 2. Bodyweight gain, total feed intake and FCR in various treatment groups were statistically significant (P<0.05). Body weight gain in broilers receiving formulated feed with 10% BW + 5% (w/v) HW powder was statistically significant (P<0.05) than other HW and BW supplemented diets. For the better understanding body weight gain and FCR have been shown in figure 5–6 throughout the experimental period, respectively.

**Table 2:** Growth performances of broiler chickens fed various experimental diets for 30 days

| Parameters | Treatments |  |  |  |  |
| --- | --- | --- | --- | --- | --- |
|  | Commercial<br>Control (T1) | FF+10% BW (T2) | FF+10% BW +<br>5% HW(T3) | FF+10% BW +<br>10% HW(T4) | FF+10% BW +<br>15% HW(T5) |
| Initial body weight at<br>10<sup>th</sup> day (g) | 272.09±0.04<sup>a</sup> | 272.03 ±0.08<sup>a</sup> | 272.12 ±0.03<sup>a</sup> | 273.26 ±0.71<sup>a</sup> | 272.18±0.56<sup>a</sup> |
| Final body weight at<br>30<sup>th</sup> day (g) | 1876.0±53.25<sup>b</sup> | 1628.0±123.67<sup>b</sup> | 1700.0±60.08<sup>b</sup> | 1632.0±51.23<sup>ab</sup> | 1650.0±21.21<sup>a</sup> |
| *Body weight gain (g) | 1603.91±53.25<sup>b</sup> | 1355.97±123.68<sup>b</sup> | 1427.88±60.09<sup>b</sup> | 1358.74±50.55<sup>ab</sup> | 1377.81±21.0<sup>a</sup> |
| Total feed intake (g) | 2706.80±13.37<sup>b</sup> | 2688.60±17.06<sup>a</sup> | 2619.20±12.77<sup>a</sup> | 2692.40±16.09<sup>a</sup> | 2676.20±8.71<sup>a</sup> |
| **FCR | 1.69±.06<sup>a</sup> | 2.07±.25<sup>a</sup> | 1.85±.09<sup>a</sup> | 1.97±0.08<sup>a</sup> | 1.96±.03<sup>a</sup> |
| ***Mortality (%) | 6.64±4.06<sup>ab</sup> | 19.92±6.21<sup>a</sup> | 9.96±4.06<sup>ab</sup> | 9.96±4.06<sup>ab</sup> | 3.32 ± 3.32<sup>b</sup> |

Values are expressed as mean standard error of at least five replications each of which contains six birds. Different letter within a row differ significantly ( $p < 0.05$ ). Same letters within a row are statistically non-significant ( $p < 0.05$ ).
FF=Formulated feed; BW=Buckwheat; HW=Honeyweed;
\*Body Weight gain (g) = Final body weight at 30th day –Initial body weight at 10th day.
\*\* FCR (Feed Conversion Ratio) = Total feed intake (g)/Body weight gain (g).
\*\*\*Mortality (%)=Number of dead birds / Total no of birds × 100%

### Lipid profile

The effects of formulated diets on some serum biochemical parameters like total cholesterol, HDL-cholesterol, LDL-cholesterol, triacylglycerols of broilers are shown in figure 1–4. Serum total cholesterol fed BW and HW decreased (P<0.05) compared with the control groups. It was observed that the lowest value was recorded in birds fed with 10% BW+15% (w/v) HW powder (Figure-1). HDL-cholesterol concentrations were significantly higher (P <0.05) in birds receiving diets with BW and HW (Figure-2). The highest HDL-cholesterol content was found in 10% BW+15% (w/v) HW powder at 30^th^ day. Serum LDL and triglycerides level were also decreased in the diets of BW and HW compared with the control (Figure-3, 4). The lowest LDL and triglyceride value (46.21 mg/dl), (50.61mg/dl) was recorded in birds fed 10% BW+15% (w/v).

**Figure 1:**
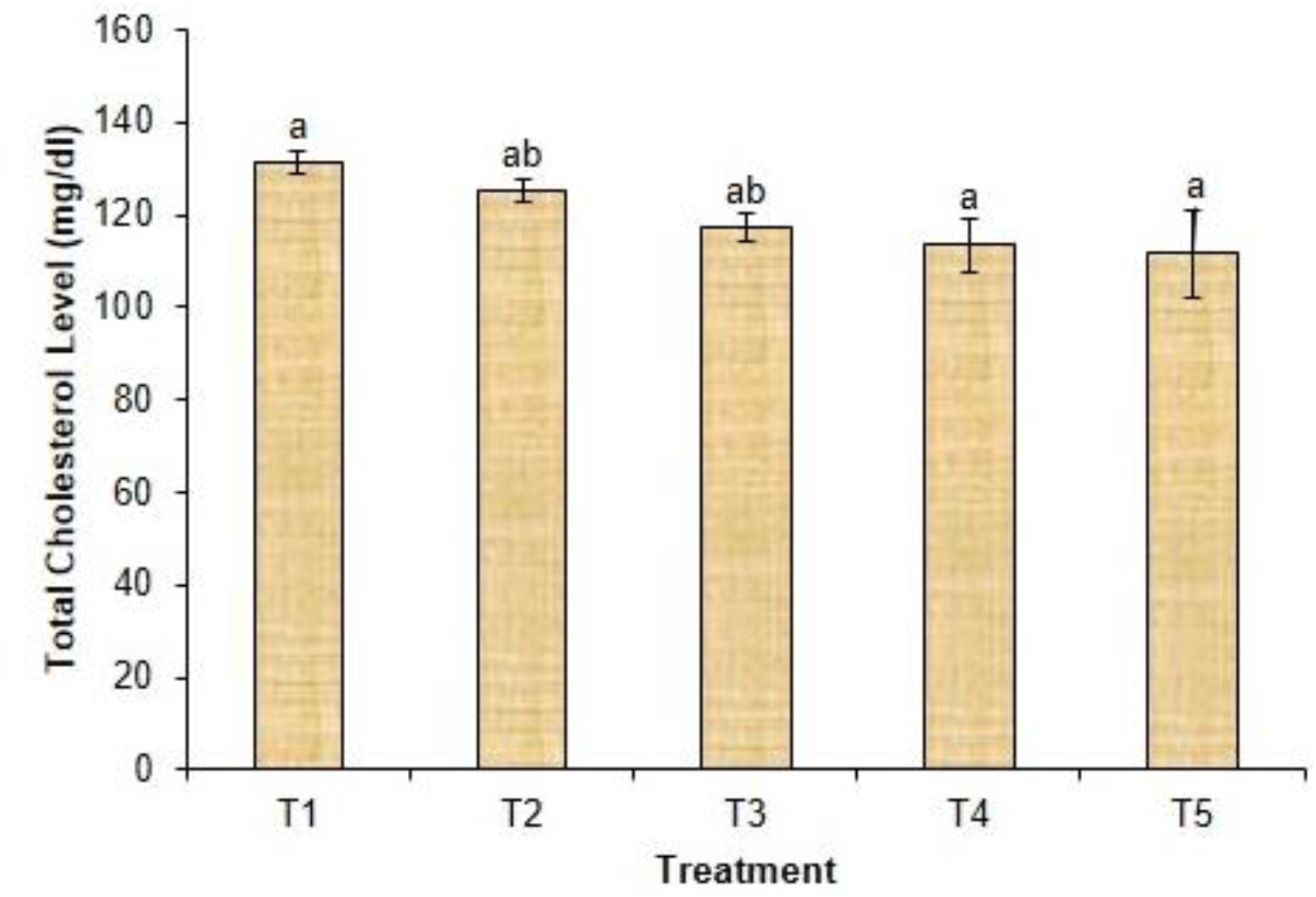
Serum total cholesterol of broiler chicks fed with commercial control (T1), Formulated feed (T2) + 10% Buckwheat, Formulated feed+10% Buckwheat+5% Honeyweed (T3), Formulated feed+10% Buckwheat+ 10% Honeyweed (T4), Formulated feed+10% Buckwheat+15% Honeyweed at 30th day of treatment. The results are expressed as mean± SD of six birds. Data point bearing different letters are significantly different at p < 0.05.

**Figure 2:**
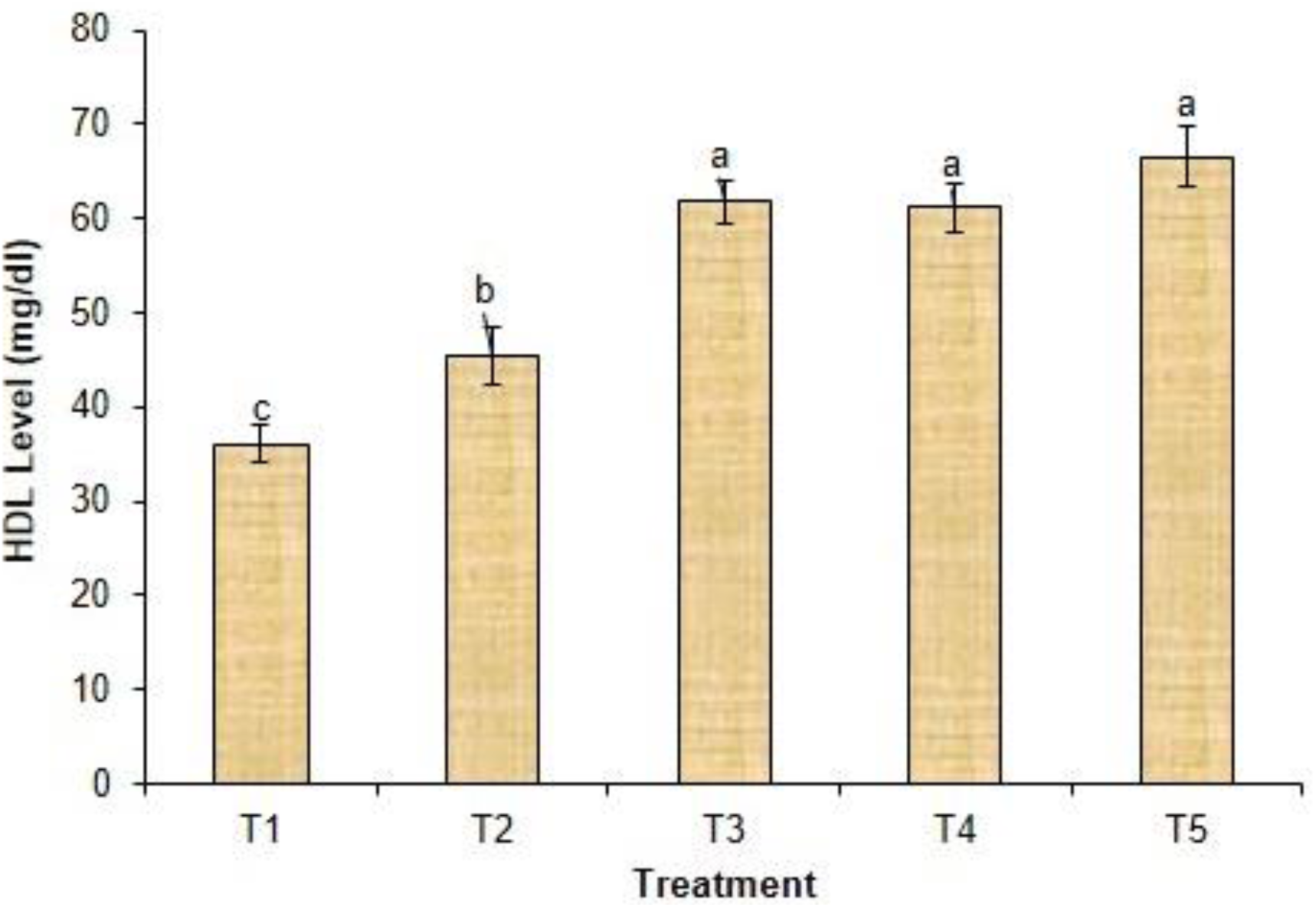
HDL level of broiler chicks fed with commercial control (T1), Formulated feed (T2) + 10% Buckwheat, Formulated feed+10% Buckwheat+5% Honeyweed (T3), Formulated feed+10% Buckwheat+ 10% Honeyweed (T4), Formulated feed+10% Buckwheat+15% Honeyweed at 30th day of treatment. The results are expressed as mean± SD of six birds. Data point bearing different letters are significantly different at p < 0.05.

**Figure 3:**
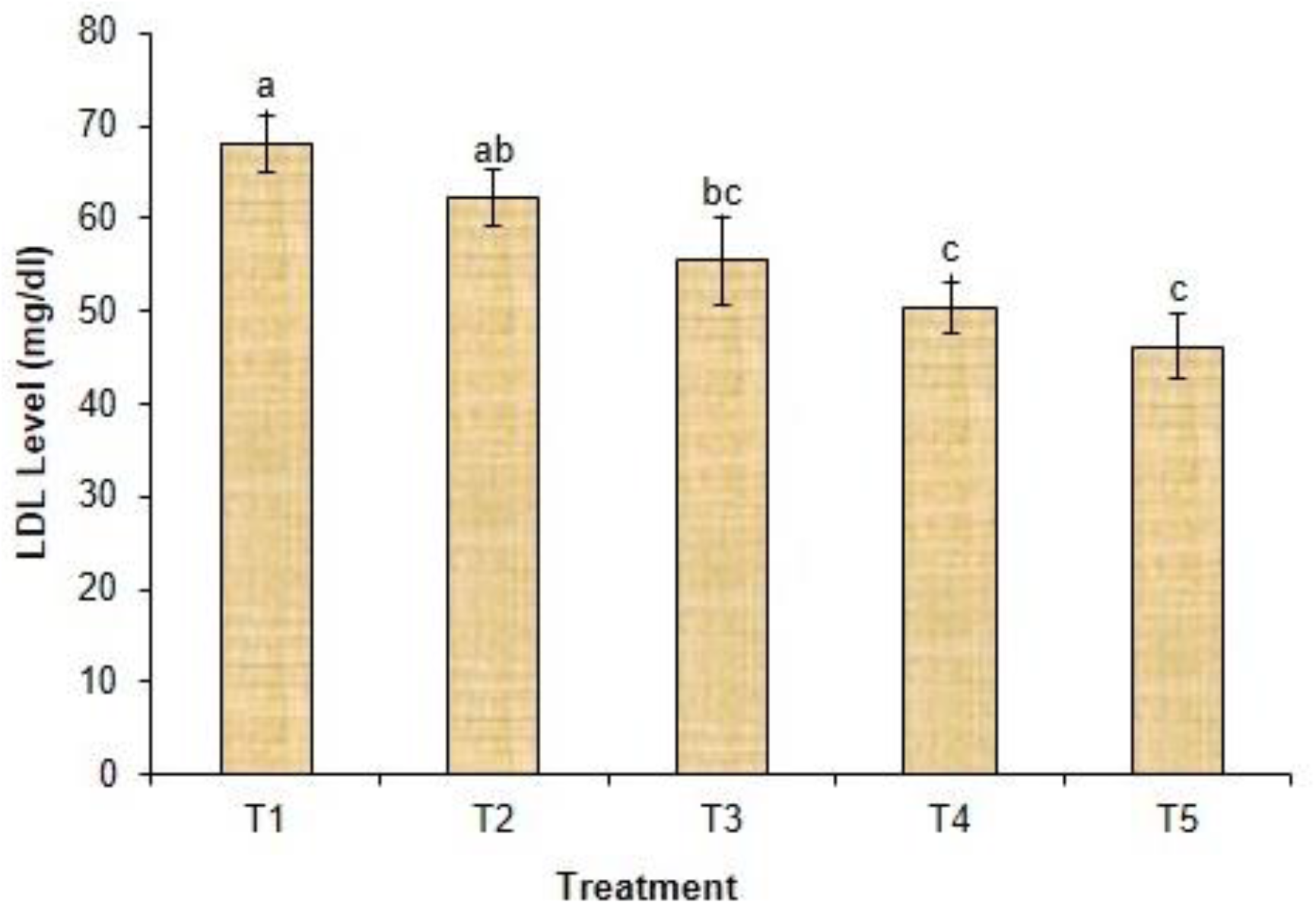
LDL level of broiler chicks fed with commercial control (T1), Formulated feed (T2) + 10% Buckwheat, Formulated feed+10% Buckwheat+5% Honeyweed (T3), Formulated feed+10% Buckwheat+ 10% Honeyweed (T4), Formulated feed+10% Buckwheat+15% Honeyweed at 30th day of treatment. The results are expressed as mean± SD of six birds. Data point bearing different letters are significantly different at p < 0.05.

**Figure 4:**
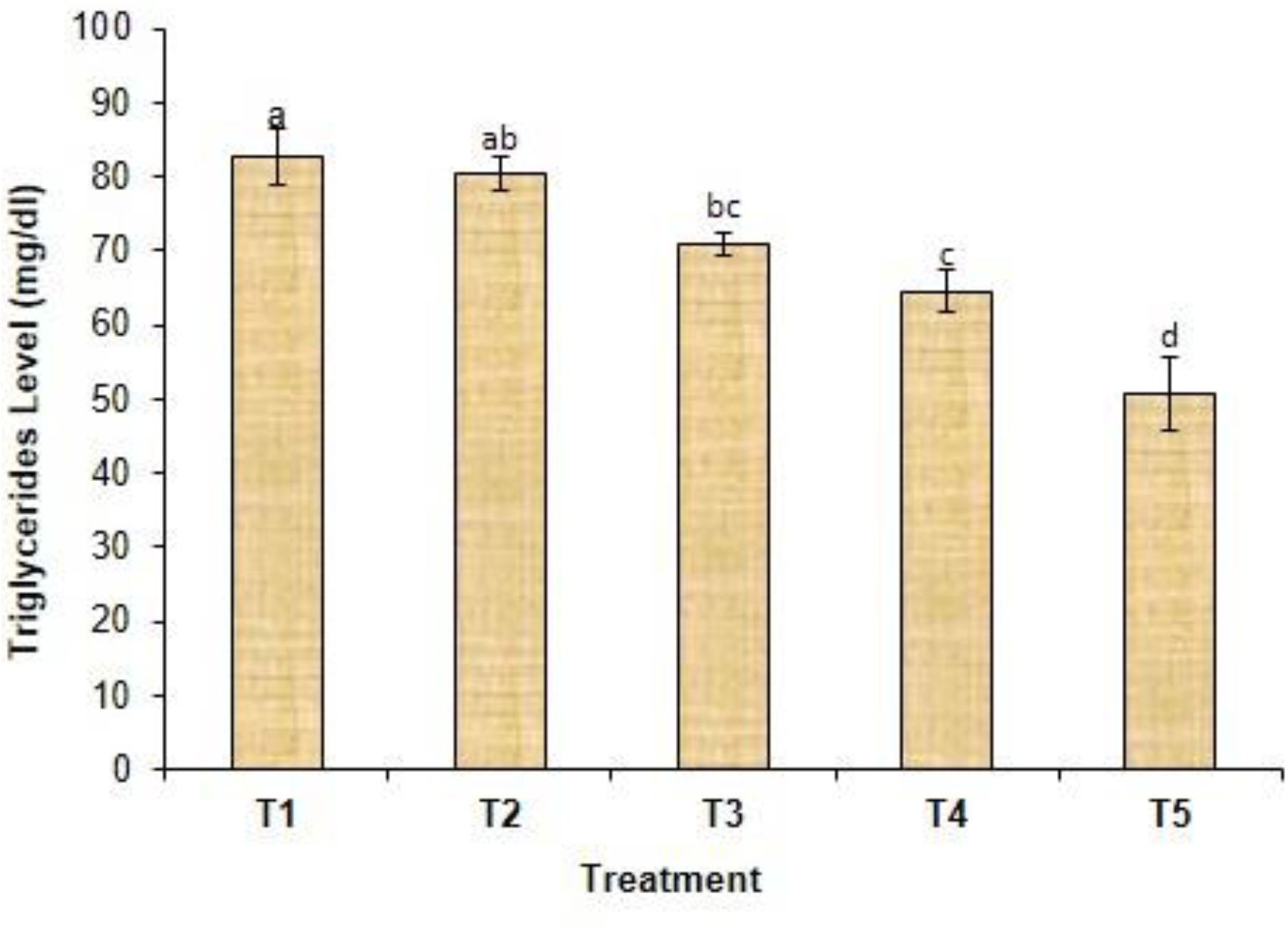
Triglyceride level of broiler chicks fed with commercial control (T1), Formulated feed (T2) + 10% Buckwheat, Formulated feed+10% Buckwheat+5% Honeyweed (T3), Formulated feed+10% Buckwheat+ 10% Honeyweed (T4), Formulated feed+10% Buckwheat+15% Honeyweed at 30th day of treatment. The results are expressed as mean± SD of six birds. Data point bearing different letters are significantly different at p < 0.05.

**Figure 5:**
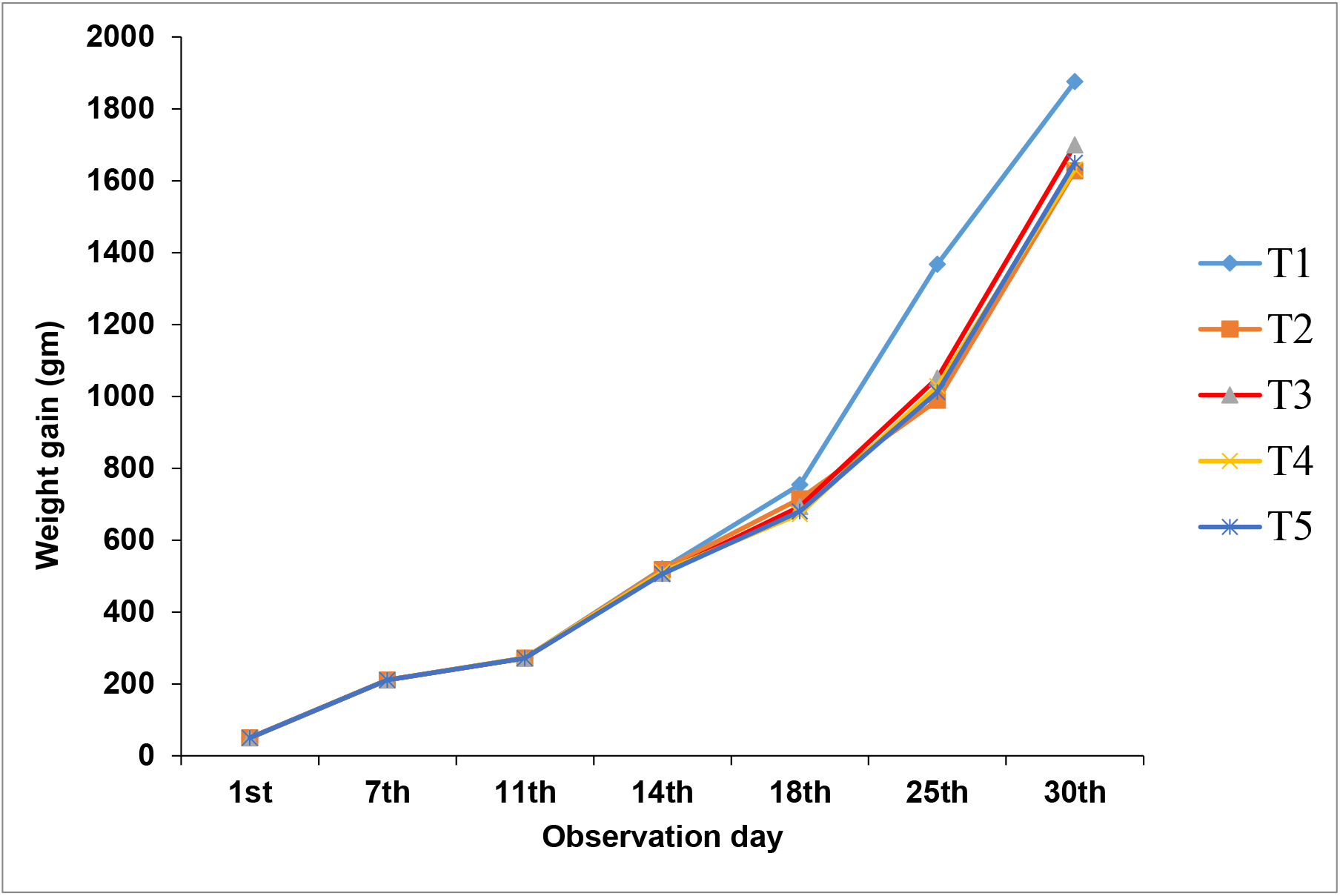
Body weight gain of broiler chicks fed with commercial control (T1), Formulated feed (T2) + 10% Buckwheat, Formulated feed+10% Buckwheat+5% Honeyweed (T3), Formulated feed+10% Buckwheat+ 10% Honeyweed (T4), Formulated feed+10% Buckwheat+15% Honeyweed at 30th day of treatment. The results are expressed as mean± SD of six birds. Data point bearing different letters are significantly different at p < 0.05.

**Figure 6:**
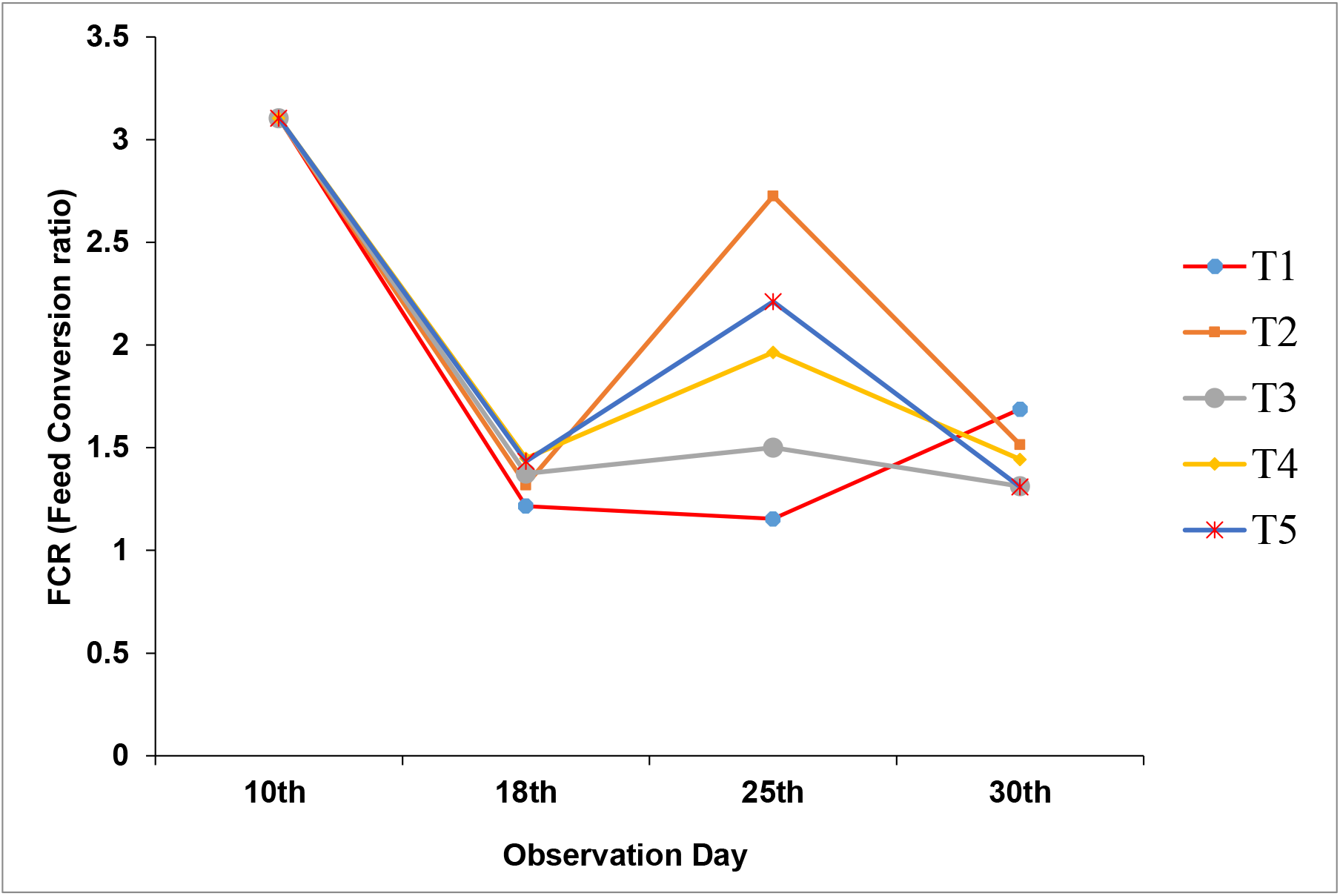
Feed Conversion Ratio (FCR) of broiler chicks fed with commercial control (T1), Formulated feed (T2) + 10% Buckwheat, Formulated feed+10% Buckwheat+5% Honeyweed (T3), Formulated feed+10% Buckwheat+ 10% Honeyweed (T4), Formulated feed+10% Buckwheat+15% Honeyweed at 30th day of treatment. The results are expressed as mean± SD of six birds. Data point bearing different letters are significantly different at p < 0.05.

### Hematological Parameters

The results of the hematological analysis of the experimental birds are present in Table 3. It was observed that there were significant (p<0.05) differences among the treatment groups in hematological parameters except the total red blood cell, mean cell volume, mean cell hemoglobin concentration (MCHC), monocytes, basophils, platelet distribution and width, hematocrit or PCV and total platelet count. The highest hemoglobin value was found (7.83 g/dl) in 10% BW+5% (w/v) HW powder (T3) and it was significantly (P<0.05) higher than the control groups. On the 30^th^ day the values of TWBC (cells 10^3^) were no significance difference (P <0.05) among the BW and HW supplement treatments groups. The highest value (7.54) was obtained in commercial control. In case of MCH there was significant difference between the control and BW and HW supplemented diet. At 30^th^ day of the experiment, the MPV parameter had significant difference (P<0.05) between control and different treatments of BW and HW. In our experimental results highest value (9.58 fl) at 10% BW+5% (w/v) HW powder (T3) and lowest value (7.23 fl) at commercial control (T1) for MPV. From table 3 it was showed that, neutrophils percentage in commercial control group showed highest result (39.23%) and the values were significantly decreased with the increase of HW percentage compared with the control groups. Lymphocytes (%) of the present study showed that there were significance difference (P<0.05) among the control group and other treatment groups. The highest result (71.31%) was obtained at commercial control (T1) and the lowest result (42.85%) was obtained at FF+10% BW (T2). Moreover, it was also found from table 3 at 30^th^ day of our experiment the results of RDW-CV (%) and P–LCR % were not statistically similar among the treatments. The statistically lowest P-LCR % was found in commercial control (T1) (29.45) and the highest % of P-LCR was found at 10% BW+5% (w/v) HW powder (T3) (33.46) compared with others treatments. It was also obsereved that the P-LCR % value significantly increased in the treatment groups due to the combined phytogenic effect of BW and HW in the supplementary diets. There were significant impact of HW and on the percentage of PCT. The highest value (0.03%) was obtained for the supplementation of 5% HW with 10% BW as diets for broilers at T3 and this value was significantly differences (P <0.05) compared with commercial control at T1.

**Table 3:** Hematological Parameters of broilers chickens fed with different experimental diets for a period of 30 days

| Parameters | Treatments |  |  |  |  |
| --- | --- | --- | --- | --- | --- |
|  | Commercial<br>Control (T<sub>1</sub>) | FF+10% BW (T<sub>2</sub>) | FF+10% BW + 5%<br>HW(T<sub>3</sub>) | FF+10% BW +<br>10% HW(T<sub>4</sub>) | FF+10% BW +<br>15% HW(T<sub>5</sub>) |
| Haemoglobin (g/dl) | 5.36±0.10<sup>b</sup> | 6.99±0.81<sup>a</sup> | 7.83±1.18<sup>a</sup> | 7.52±0.63<sup>a</sup> | 7.10±0.59<sup>a</sup> |
| Total WBC (cells 10<sup>3</sup>/μl) | 7.54±3.09<sup>a</sup> | 6.14±2.44<sup>ab</sup> | 3.37±0.91<sup>b</sup> | 3.38±0.70<sup>b</sup> | 4.50±1.80<sup>b</sup> |
| Total RBC Count (cells<br>10<sup>6</sup>/μl) | 2.88±0.44<sup>a</sup> | 3.41±0.29<sup>a</sup> | 3.47±1.21<sup>a</sup> | 3.37±0.62<sup>a</sup> | 3.43±0.66<sup>a</sup> |
| MCV (fl) | 133.1±6.34<sup>a</sup> | 132±9.41<sup>a</sup> | 136.27±0.65<sup>a</sup> | 133.38±4.87<sup>a</sup> | 137.82±1.92<sup>a</sup> |
| MCH (pg) | 19.50±0.13<sup>b</sup> | 26.49±0.52<sup>a</sup> | 26.94±0.03<sup>a</sup> | 26.25±0.03<sup>a</sup> | 25.77±2.89<sup>a</sup> |
| MCHC (g/dl) | 23.56±7.04<sup>a</sup> | 26.49±0.52<sup>a</sup> | 25.78±5.62<sup>a</sup> | 27.55±3.62<sup>a</sup> | 22.96±3.06<sup>a</sup> |
| MPV (fl) | 7.23±0.04<sup>b</sup> | 9.37±0.76<sup>a</sup> | 9.58±0.88<sup>a</sup> | 9.40±1.12<sup>a</sup> | 9.14±0.69<sup>a</sup> |
| PDW (fl) | 11.05±2.10<sup>a</sup> | 12.80±0.62<sup>a</sup> | 12.29±0.28<sup>a</sup> | 12.24±2.03<sup>a</sup> | 12.20±2.36<sup>a</sup> |
| Neutrophils (%) | 39.23±1.09<sup>a</sup> | 34.09±2.68<sup>b</sup> | 35.20±1.30<sup>b</sup> | 36.51±3.86<sup>ab</sup> | 34.60±.54<sup>b</sup> |
| Lymphocytes (%) | 71.31±.51<sup>a</sup> | 42.85±2.04<sup>b</sup> | 46.72±1.22<sup>b</sup> | 45.80±3.27<sup>b</sup> | 44.66±6.28<sup>b</sup> |
| Monocytes (%) | 4.59±0.53<sup>a</sup> | 4.42±0.72<sup>a</sup> | 4.38±0.61<sup>a</sup> | 4.01±0.25<sup>a</sup> | 4.21±0.51<sup>a</sup> |
| Basophils (%) | 1.43±0.06<sup>a</sup> | 1.27±0.42<sup>a</sup> | 1.37±0.41<sup>a</sup> | 1.20±0.44<sup>a</sup> | 1.28±0.47<sup>a</sup> |
| Hematocrit or PCV (%) | 38.21±0.42<sup>a</sup> | 38.84±2.76<sup>a</sup> | 39.67±4.47<sup>a</sup> | 38.63±10.02<sup>a</sup> | 39.08±2.48<sup>a</sup> |
| RDW –CV (%) | 8.57±0.19<sup>b</sup> | 8.76±0.17<sup>b</sup> | 9.34±0.17<sup>a</sup> | 8.62±0.28<sup>b</sup> | 8.74±0.33<sup>b</sup> |
| P-LCR (%) | 29.45±5.01<sup>b</sup> | 32.36±2.51<sup>ab</sup> | 33.46±0.90<sup>a</sup> | 32.47±0.19<sup>ab</sup> | 33.18±1.06<sup>a</sup> |
| Total Platelet Count (per<br>cmm) | 170.99±10.46<sup>a</sup> | 170.94±13.30<sup>a</sup> | 177.40±9.26<sup>a</sup> | 174.44±3.12<sup>a</sup> | 175.51±7.20<sup>a</sup> |
| PCT (Plateletcrit) (%) | 0.00±0.00<sup>d</sup> | 0.01±0.00<sup>c</sup> | 0.03±0.00<sup>a</sup> | 0.02±0.00<sup>b</sup> | 0.02±0.00<sup>b</sup> |

Values are expressed as mean standard deviation of at least five replications each of which contains six birds. Different letter within a row differ significantly ( $p < 0.05$ ). HW= Honey Weed; OC= Own control; WBC= White blood cell; RBC= Red blood cell; PCV=Packed cell volume; MCV= Mean cell volume; MCH=Mean cell haemoglobin; MCHC=Mean cell haemoglobin concentration; P-CLR=Platelet large cell ratio; RDW-CV= Red cell distribution and Width coefficient variation; PCT=Plateletcrit; PDW=Platelet distribution and Width; MPV= Mean platelet volume.

## Discussion

In current findings, it is proved that there is no inferior issues on the growth performance and feed intake of broilers fed with only BW or in combination with HW extracts. This report first time indicates that feeding BW and HW extracts positively favours the lipid profiles and blood parameters of broilers. Our current findings, recommended that phytogenic feed additives BW and HW extracts might be an alternative solution to synthetic antibiotics. In various study, it has been noticed that BW has the potentiality to use as an alternative to antibiotics due to the presence of pharmacologically active constituents such as flavones, flavonoids, phytosterols, tocopherols, inositol phosphates, rutin and myo-inositol (Zhang *et al*., 2012). Pharmacological compound rutin is the most prominent in buckwheat, and is reported to improve immunity (Sayed *et al*., 2015). Leonurine is the most influential pharmacological compound in honeyweed that significantly protect cardiovascular diseases (Zhang *et al*., 2018). The poor performance of chickens fed higher levels of BW and HW was probably may be due to the excess amount of HW can create taste variation in feed. In this current findings, feed conversation ratio (FCR) is better than previously experimented combination of buckwheat and black cumin (Islam *et al*., 2016), buckwheat and chitosan (Sayed *et al*., 2015) or only buckwheat supplemented diet (Sayed *et al*., 2013).

According to the results on the lipid profile of broilers in this study it could be deduced that BW and HW have favourable effects on serum metabolites. BW has been used for centuries for medicinal purposes and reported to decrease serum lipid profile. Supplementation of BW powder in broiler diet significantly (p <0.05) decreased serum triglycerides but elevated the HDL contents in broiler (Sayed *et al*., 2013). On the other hand, plasma cholesterol decreased and high density lipoprotein-cholesterol increased has been reported in *L. sibiricus* herb extract supplemented feed in mice (Lee *et al*.,2010). Triglyceride (TG) accumulation inhibitory effects has been reported in free fatty acid-induced HepG2 cells due to flavonoids leonurusoides (Zhang *et al*., 2013) in HW. Our current findings is more or less similar to buckwheat and black cumin supplemented feed (Islam *et al*., 2016). Triglyceride level also show good result than only buckwheat supplemented feed in broilers (Sayed *et al*., 2013). Thus BW with small amount of HW extract in broiler diets could be a potential alternative for synthetic feed additives in broilers.

Hemoglobin is an iron containing protein which transports oxygen from the lung to the body. It was observed that, the hemoglobin value of BW and HW supplemented diet higher than the control groups and the percentage of hemoglobin was decreased from medium and high dose of *Leonurus sibiricus* in rats (Chua *et al*., 2008). Intriguingly, our study showed that BW and HW supplement diet increases hemoglobin. White blood cells are the part of immune system. At 10% BW+5% HW (T3) obtained result was closer to the normal range which may mean that the antioxidative property of BW and HW influenced the immune system of response in broiler chicken. The major function of the red blood cells is to transport hemoglobin, which in turn carries oxygen from the lungs to the tissues. Present value of TRBC in combined BW and HW supplemented feed is slightly higher than normal range in broiler studied by (Orawan, 2007). In case of Mean cell volume there were not significance differences (P<0.05) among the all groups. It has been reported that MCV was decreased in the treated groups with higher concentration of flavonoid compounds in rats (Fausto *et al*., 2017). Interestingly, Mean Cell Hemoglobin (MCH) and MCHC were more or less similar that has been observed in the ginger root powder supplemented diets (Zomrawi *et al*., 2013) on broiler chicks. MPV and PDW increases the platelet production. Moringa leaves inclusion diet in broiler improve the MPV and PDW (Zomrawi *et al*., 2013). Neutrophils are a type of white blood cell that helps to heal damaged tissues and resolve infections. Neutrophil blood levels increase naturally in response to infections, injuries, and other types of stress. They may decrease in response to severe or chronic infections, drug treatments, and genetic conditions. Neutrophils percentage is decreased significantly in BW and HW supplemented feed compared with commercial control group that are the normal range of neutrophils was found 15-40% for chickens (Jain 1993). In case of fermented garlic powder supplementation feed lymphocytes percentage is increased with the increase of garlic powder application (Ao *et al*., 2011) but in present study lymphocytes percentage is decreased with the increase of HW percentage. There was no significance changed of basophils percentage in broiler chickens for the phytogenic combined effect of BW and HW and the values are almost similar of normal basophil percentage in broilers (Orawan, 2007). PCV (%) value provides information about how much blood is comprised of red blood cells. A low score on the range scale may be a sign that have too little iron, the mineral that helps to produce red blood cells. A high score could mean dehydrated or have another condition. Similar PCV % was observed by (Oke *et al*., 2017) for supplementary diet of olive leaf extract in broiler. The P-LCR % value significantly increased in the treatment groups due to the combined phytogenic effect of BW and HW in the supplementary diets for broiler chickens but our results higher than previously studied Olive oil, Black seed oil, Flax seed oil supplemented diet (Ashraf *et al*., 2019).

## Conclusion

Based on the findings it may be concluded that combination of BW and HW supplemented diets have positive impact on broiler growth rate. Diet containing 10% BW with 5% HW (T3) may be used in poultry industry for increasing the rate of poultry as it showed highest body weight gain and lowest FCR rate compared to other BW and HW supplemented diet. Moreover, the 10% BW with 5% HW (T3) treatment was found to have good impact on lowering the health hazardous serum total cholesterol, LDL, triacylglycerols along with elevation of serum HDL-cholesterol level and improves blood parameters than commercial control. Thus, poultry industries can produce safe and low cholesterol poultry meat as there is a positive correlation between serum cholesterol content and meat cholesterol. However, further in-depth research is highly recommended for assessing the depth mechanisms of blood parameters and lipid profile lowering effect of BW and HW in broiler chickens.

## Acknowledgements

We are grateful to Ministry of Science and Technology, Government of the Peoples’ Republic of Bangladesh for funding this research work. We are also grateful to Institute of Research and Training (IRT), Hajee Mohammad Danesh Science and Technology University, Dinajpur, Bangladesh for their financial support to complete the study.

